# Serum and sputum MMP-9/TIMP-1 in winter sports athletes and swimmers: relationships with airway function

**DOI:** 10.1101/2021.02.10.430578

**Authors:** Valérie Bougault, Julie Turmel, Louis-Philippe Boulet

## Abstract

**Introduction:** Cross-country skiers and swimmers present characteristics of airway inflammation and remodeling of the extracellular matrix similar to what is observed in mild asthma. We aimed to compare serum and sputum MMP-9/TIMP-1 levels, to assess the balance between airway fibrogenesis and inflammation process in both categories of athletes, and to observe its seasonal variations in winter sports athletes.

**Methods:** Competitive winter sports athletes (n=41), swimmers (n=25) and healthy nonathletes (n=10) had blood sampling, lung function measurement, skin prick tests, eucapnic voluntary hyperpnea challenge, methacholine inhalation test (MIT), and induced sputum analysis. Twelve winter sport athletes performed the test during both summer and winter. Serum and sputum MMP-9 and TIMP-1 were measured by ELISA.

**Results:** No significant difference in serum or sputum MMP-9/TIMP-1 ratio was observed between groups, nor relationship with airway function or responsiveness. Serum MMP-9/TIMP-1 ratio was significantly higher during the summer in winter sport athletes compared with winter season (median [Interquartile range]: 3.65 [2.47-4.03] ng.ml^-1^ and 1.27 [0.97-1.62] ng.ml^-1^, respectively, p=0.005). Sputum MMP-9 correlated with Methacholine PC_20_ and serum CC16/SP-D ratio.

**Conclusion:** MMP-9/TIMP-1 ratio in sputum or serum may fluctuate with training or environment but does not correlate with airway lung function or responsiveness in competitive athletes.

## Introduction

Competitive swimmers and winter sports athletes are particularly at risk to develop respiratory disease or symptoms during their career, due to the high levels of ventilation sustained during prolonged training periods in very specific environmental conditions (i.e. inhalation of cold dry air in winter sports athletes and chlorine by-products in swimmers) (Karjalainen *et al*. 2000, Bougault *et al*. 2010, Montjoy *et al*. 2015). On one hand, winter sports athletes frequently report exercise-induced respiratory symptoms, especially cough, due to cold dry air inhalation and disturbing their training or performance, but not necessarily related to exercise-induced bronchoconstriction (EIB) or airway hyperresponsiveness (AHR) (Bougault *et al*. 2010, Stenfors *et al*. 2010, Rundell *et al*. 2015, Boulet *et al*. 2017). On the other hand, a majority of competitive swimmers present with AHR to direct or indirect challenge in a Lab, without complaining of respiratory symptoms or having EIB during swimming, except maybe when chlorine byproducts are elevated in the atmosphere of swimming pools (Potts 1994, Levesque *et al*. 2006, Bougault *et al*. 2010, Couillard *et al*. 2014). Both categories of competitive athletes’ present characteristics of airway remodeling of the extracellular matrix (ECM), such as fibrosis of the submucosa and reticular basement membrane thickening, whatever the AHR status to methacholine (Karjalainen *et al*. 2000, Bougault *et al*. 2012). The inflammatory profile in airway tissue looks, however, different in competitive cross-country skiers and swimmers, characterized by a marked increase in T-lymphocytes and macrophage cell counts in the former and mast cells infiltration in the latter (Karjalainen *et al*. 2000, Bougault *et al*. 2012). All these observations may indicate a different airway modification phenotype according to the training environment.

In some phenotypes of asthma, an imbalance between the expression of proteases and anti-proteases within the ECM, and particularly of matrix metalloproteinase-9 (MMP-9) and its inhibitor (TIMP-1), is believed to be a biomarker of tissue destruction or abnormal repair, and fibrosis (Mautino *et al*. 1997, 1999a, 1999b, Tanaka *et al*. 2000, Gueders *et al*. 2006). The activation of epithelial cells, and the recruitment of neutrophil and mast cells (Nanson *et al*. 2001, Boulet *et al*. 2005, Chimenti *et al*. 2010, Kippelen *et al*. 2010, Hallstrand *et al*. 2013), as observed during exercise ventilation, may stimulate the secretion or release of matrix metalloproteinase-9 (MMP-9) (Mattos *et al*. 2002, Cundall *et al*. 2003 Barbaro *et al*. 2014 Ventura *et al*. 2014). In excess, MMP-9 and tissue-inhibitor (TIMP-1) have been shown to lead to airway tissue damage and fibrosis in asthma (Mautino *et al*. 1997, 1999a, 1999b, Gueders *et al*. 2006, Grzela *et al*. 2016). In order to distinguish between both potential phenotypes, we compared the ratio of MMP-9 and its inhibitor (TIMP-1) during the annual beginning of the new training season in winter sport athletes and swimmers, in relation with other classical biomarkers of inflammation. Whereas a low MMP-9/TIMP-1 ratio would rather reflect a predominant fibrogenesis over airway inflammation, an increased ratio would indicate a predominant inflammatory process, which seems also to predict a better response to inhaled corticosteroids in asthma (Bosse *et al*. 1999). The secondary objective was to observe the variations of MMP-9/TIMP-1 ratio between winter and summer, in winter sports athletes only, because if swimmers are exposed to the pool air all year round, winter sport athletes are subjected to climatic variations related to their sport.

## Material and methods

### Participants

Blood samples from subjects (athletes and controls) participating to previous studies on airway reactivity within 2 years (2007-2008) were retrospectively analyzed within 5 years after sampling. Inclusion criteria for inclusion of subjects were (1) having participated to a previous study with a similar protocol for all (see below for the study design), (2) having been tested during the annual recovery period or the beginning of the sport season for athlete (summer for winter sport athletes and end of summer-autumn for swimmers). Subjects had to be from 14 to 35 years of age, nonsmokers, and free of any disease that may have interfered with the study. Athletes had to be active competitors, training at least 10 hours per week. Non-athletes non-asthmatic controls taking part in physical activities more than 4 hours per week or regularly exposed to a chlorinated environment were excluded. Subjects using inhaled corticosteroids on a regular basis were also excluded. All the subjects gave their written informed consent and the study protocol was approved by the IUCPQ Institutional Ethics Committee (CER 20159 and CER 20141). Data from the lung function and responsiveness analysis between both populations were already published in a previous article (1). Written informed consent was obtained from each subject before inclusion in their respective study. Retrospective study protocols and analyses (CER20141, CER20159) were approved by our institutional Ethics Committee (FWA00003485). Studies protocols were previously registered at www.clinicaltrials.gov (NCT 00686452 and NCT 00686491).

### Study design

This cross-sectional retrospective analysis was based on data from previous and ongoing prospective studies performed at our research center. The design of the tests has been previously described (Bougault *et al*. 2010). The visits were performed during the beginning of a new training season, mainly from the end of August to early October in swimmers and non-athletes, and during the summer in winter sport athletes. To check a potential change in serum biomarkers during the seasons, 12 winter sport athletes had also a blood sample during the competitive season in winter. Each test was performed at least 14h after the last low intensity training session for all participants. A physical examination and allergy skin prick tests were done and a locally developed standardized questionnaire was administered to all the subjects regarding respiratory health and sport activities. Blood samples were taken before performing an eucapnic voluntary hyperpnea challenge (EVH), followed by a methacholine inhalation test (MIT) approximately 45 min later. Both tests were performed to characterize airway responsiveness to indirect and direct stimuli, respectively. A sputum induction was also carried out in all the subjects, about one hour after the end of EVH test. The order of the tests was the same at each visit.

### Atopy

Atopic status was determined by skin-prick tests with common airborne allergens, including animal danders, house-dust mite, mixed trees, mixed grasses, pollens, and molds. The presence of atopy was defined as at least one positive (>3 mm mean wheal diameter at 10 min) response to allergens in the presence of a negative saline control and a positive skin response to histamine.

### Lung function

Spirometry was carried out according to the American Thoracic Society (ATS) specifications (Miller *et al*. 2005). Predicted spirometric values were defined according to Knudson *et al*. (1983). Three reproducible measurements of forced expiratory volume in one second (FEV_1_) and forced vital capacity (FVC) were obtained.

### Airway responsiveness to Eucapnic Voluntary Hyperpnea and Methacholine challenge

Eucapnic voluntary hyperpnea challenge (EVH) and Methacholine inhalation test (MIT) were performed using the method described by Anderson and Brannan (2003), and the 2-minute tidal breathing method (Crapo *et al*. 2000), respectively. The methacholine PC_20_, which is the concentration of methacholine inducing a 20% fall in FEV_1_, and the post-EVH fall in FEV_1_ were used for analysis. Airway hyperresponsiveness (AHR) to EVH was defined by a FEV_1_ fall of at least 10% sustained for at least two consecutive time-points after EVH and to methacholine by a PC_20_ < 16 mg.mL^-1^.

### Induced sputum

Sputum was induced using the method described by Pizzichini *et al*. (1996) and processed within 2 h following induction. Briefly, mucus plugs were selected from saliva, weighed, treated with four times their volume of dithiothreitol (DTT) (Cal-biochem Corp, La Jolla, CA, USA) and rocked for 15 min. The reaction was stopped with an equal volume of Dulbecco’s phosphate buffered saline (D-PBS) (Life Technologies, Gibco BRL, Burlington, ON, Canada), filtered and counted to determine total cell count and viability. The samples were centrifuged and supernatants frozen at −80°C until further analysis.

### Blood sampling

Blood was sampled into gel tube with a clotting accelerator. Just after sampling, the tube was let in ice in a standing position for about 30 minutes to let the clot to form. Then, the blood was centrifuged at 4°C, 1100g for 20 minutes, and micro-tubes were stored at -80°C until analysis.

### Serum and sputum analysis

Blood level of fibrinogen and C-reactive protein (CRP) were measured at the biochemistry service of the hospital by colorimetry and turbidimetry, respectively. Serum club cell proteins (cc16) and surfactant proteins D (SP-D) were measured by ELISA (Biovendor, LLC, Candler, NC, USA), according to the manufacturer’s instructions. Concentrations were determined using a microplate reader (Molecular Devices, Microplate Reader, Sunnyvale, CA, USA). The sensitivity of the assay was 500 pg.ml^-1^ for cc16 and 2.2 ng.ml^-1^ for SP-D. The ratio of cc16/SP-D was calculated and kept for analysis. Serum and sputum-supernatant of MMP-9 and TIMP-1 measurements were also done by enzyme-linked immunosorbent assay (Amersham, Oakville, ON, Canada), according to the manufacturer’s instructions. The sensitivity of the assay was 0.156 ng.mL^-1^ for MMP-9 and 0.08 ng.mL^-1^ for TIMP-1. The ratios of MMP-9/TIMP-1 were then calculated.

### Statistical analysis

Quantitative data were expressed as mean ± SD, or median [Interquartile range]. Methacholine PC_20_ is expressed as geometrical mean [Interquartile range]. Qualitative data are expressed as the number of subjects and/or percentages. Univariate normality assumptions were verified with the Kolmogorov-Smirnov test. To compare the groups, one-way ANOVA was used for age, height, training (y/week), spirometric values, serum MMP-9, and Kruskal-Wallis test was used to compare mass, training (h/week), percentage fall in FEV_1_ post-EVH, methacholine PC_20_, serum fibrinogen, CRP, TIMP-1, cc16, SP-D, cc16/SP-D ratio, and MMP-9/TIMP-1 ratio. To compare sputum values between swimmers and winter sport athletes, a paired-t test was used for neutrophil, macrophage and lymphocyte cell counts, MMP-9, MMP-9/TIMP-1 and a Mann-Whitney test for sputum total cell count, eosinophil and bronchial cell counts, and TIMP-1. The Dunn’s or Holm-Sidak’s multiple comparisons procedure were applied post hoc to the ANOVA. To compare data between winter and summer, a paired t-test or non-parametric Wilcoxon test was used in winter sport athletes, when normality or equality of variance failed. Correlations were evaluated using Pearson’s (r_p_) or Spearman (r_s_) test and the significance using a one-sample *t* test. The software used for statistical analysis was Sigmastat (version 3.5, Systat Software GmbH, Erkrath, Germany) with a limit of significance of p < 0.05. When a p < 0.05 was observed between groups, the Cohen’s d were calculated (Effect Size).

## Results

Data from 30 swimmers, 41 winter sport athletes and 30 controls respected the inclusion criteria. However, due to a technical error with the samples, blood measurement were performed in 25 swimmers and only 10 controls. The characteristics of the remaining participants are presented in Table 1.

**Table 1.**
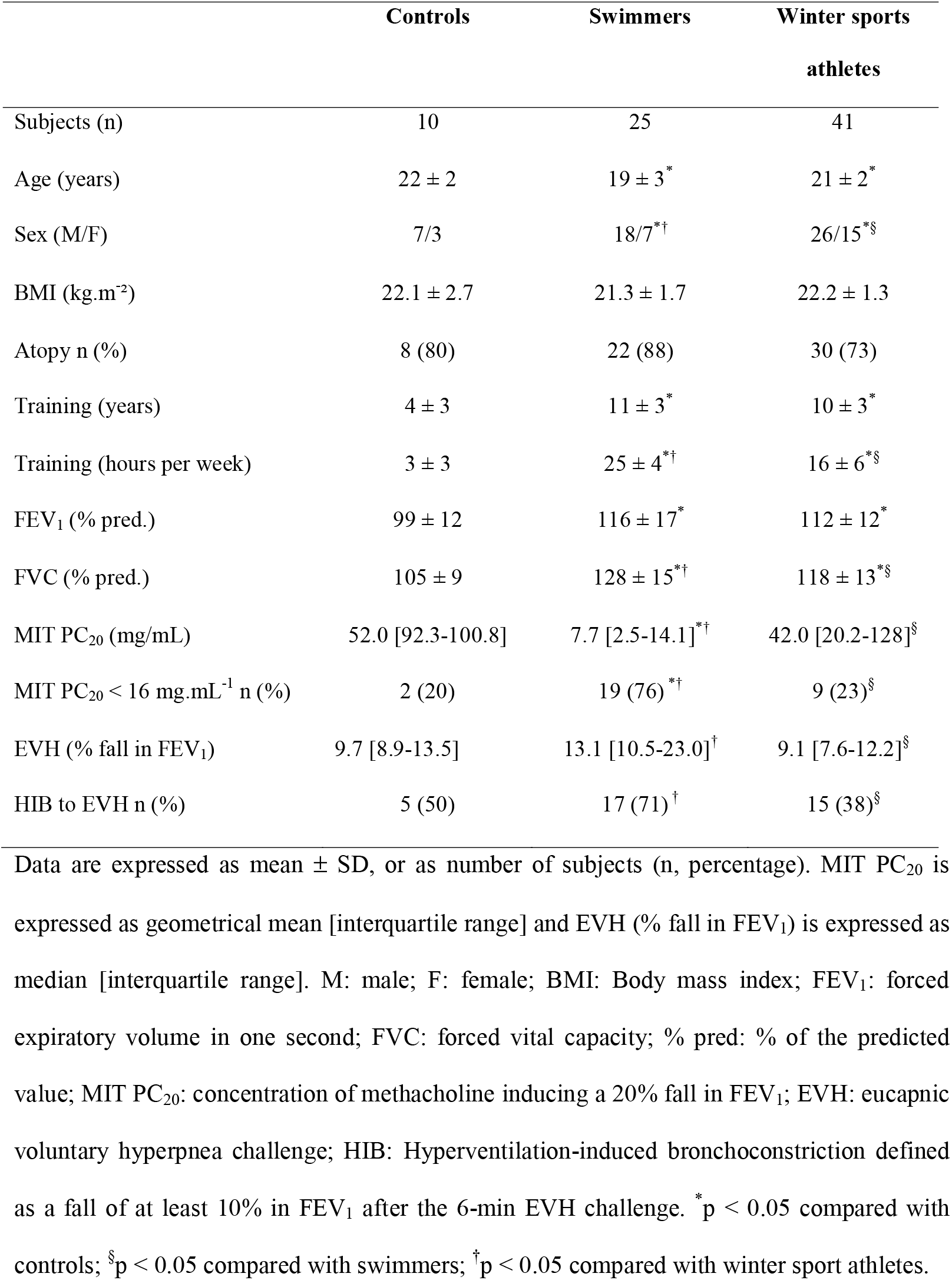
Characteristics of the participants

### Comparison of biomarkers in serum and sputum of swimmers, winter sports athletes and non-athletes

No significant difference was observed in serum CRP, fibrinogen, cc16/SP-D ratio, MMP-9, or MMP-9/TIMP-1 ratio between the three groups, during the same training season (Figure 1). The level of serum TIMP-1 was significantly higher in swimmers compared with winter sports athletes (median [interquartile range]: 208 ng.mL^-1^ [184-224] in swimmers *vs* 182 ng.mL^-1^ [162-195] in winter sports athletes, p=0.005, d=0.55), but no difference was observed between both groups of athletes and controls (median [interquartile range] in controls: 188 ng.mL^-1^ [159-196]).

**Figure 1.**
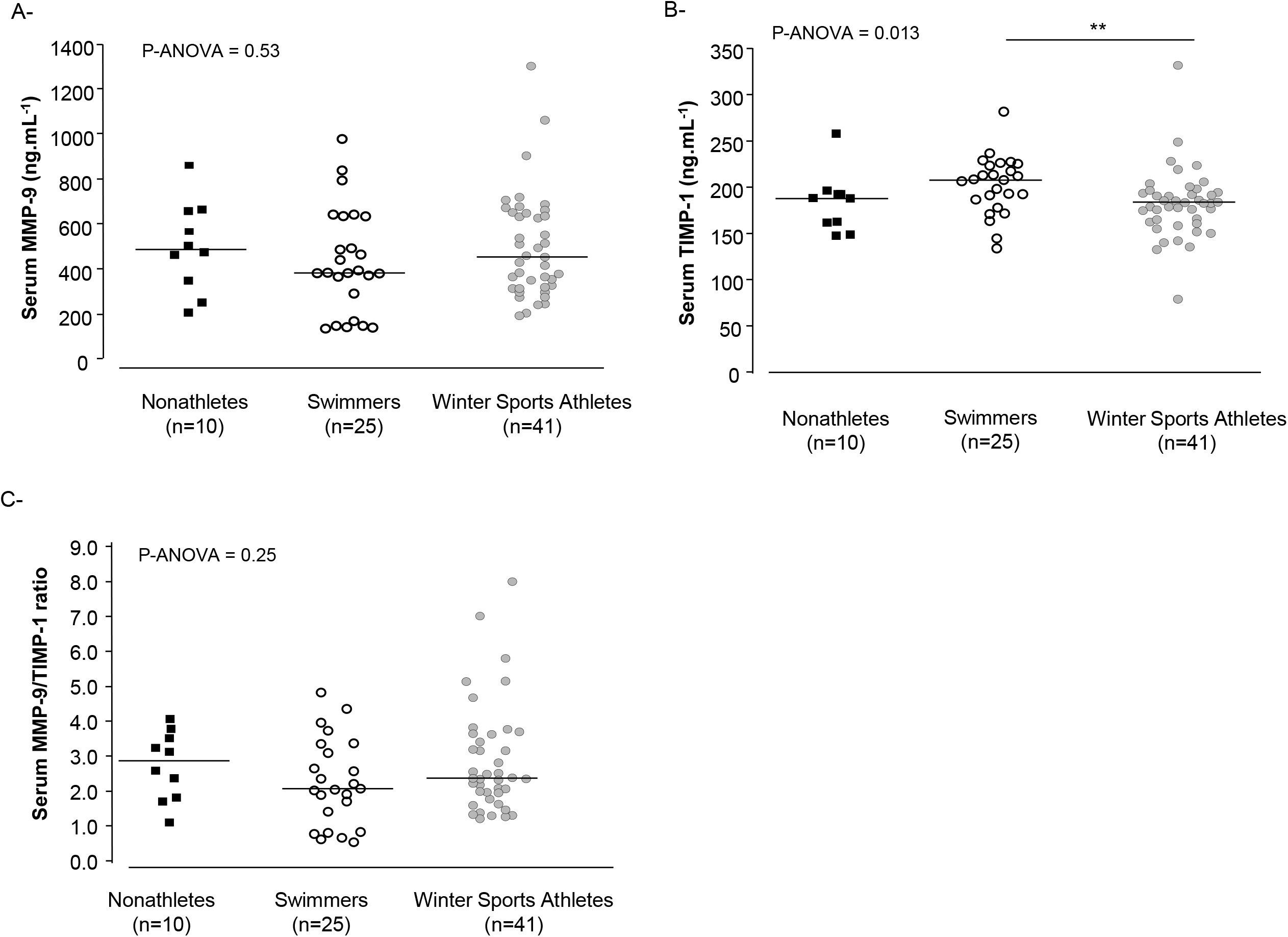
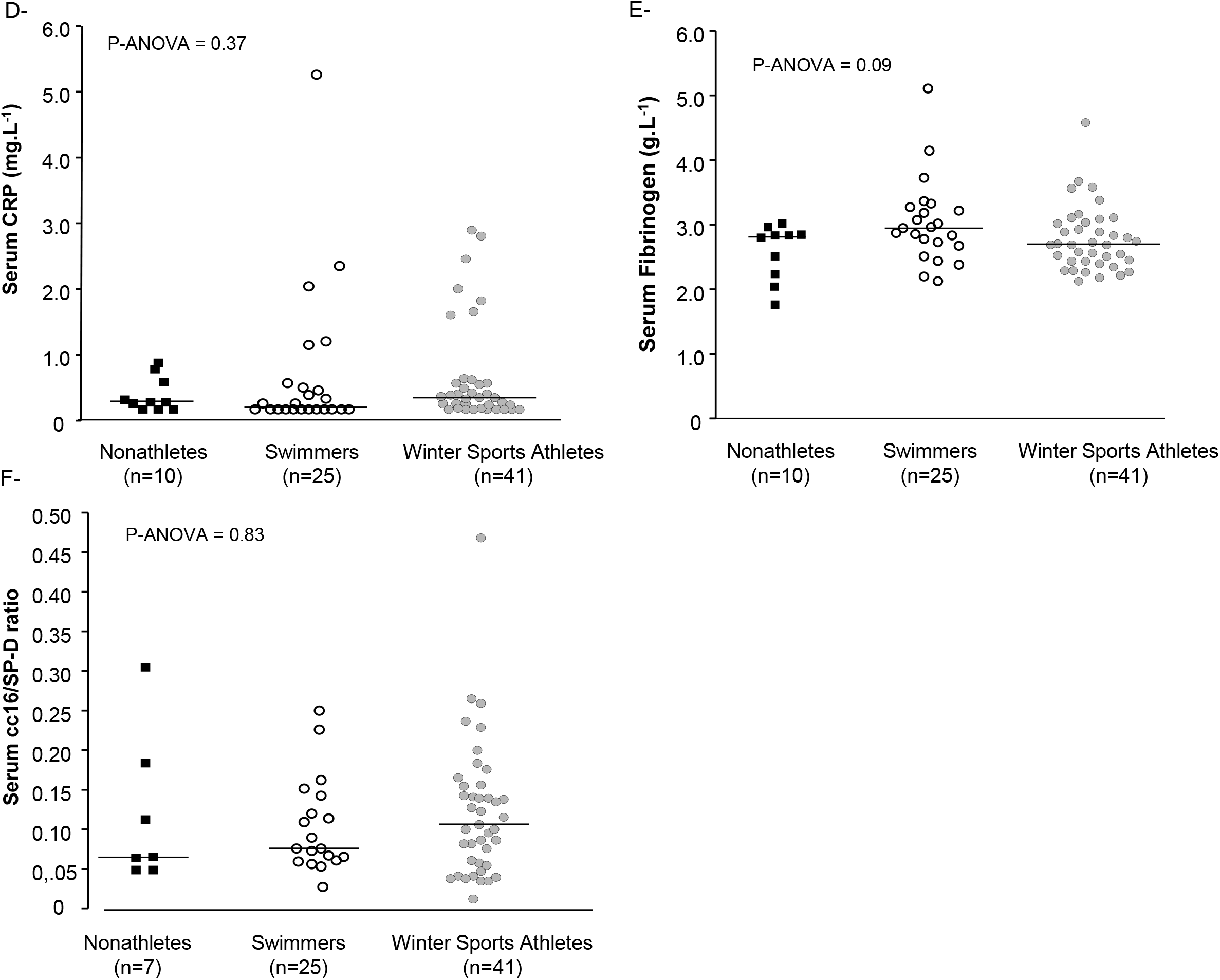
Serum values of inflammation/remodeling biomarkers in controls, swimmers and winter sports athletes. (A) MMP-9: Metalloproteinase-9; (B) TIMP-1: Tissue inhibitor of metalloproteinase-1; (C) MMP-9/TIMP-1 ratio; (D) CRP: C-reactive protein; (E) Fibrinogen; (F) cc16: Club cell proteins; SP-D: surfactant protein-D. A value of cc16/SP-D ratio of a control subject was retired from the graph but not statistical analysis, as it was an extreme data. The horizontal bar represents the median for each group. ** Post-hoc P <0.01 between groups

Sputum was obtained in 10 swimmers (40%), and 19 winter sports athletes (46%) and in only 3 controls. Only data from athletes were thus analyzed for sputum (Table 2). No significant difference was observed in sputum MMP-9, TIMP-1 or MMP-9/TIMP-1 ratio between both groups of athletes (Figure 2).

**Table 2.** Sputum cell counts characteristics in participants.

|  | Swimmers | Winter sports athletes |
| --- | --- | --- |
| Subjects (n) | 10 | 18 |
| Total cell count ($\times 10^6$ cells) | 0.485 [0.203-1.745] | 0.366 [0.143-0.986] |
| Neutrophils (%) | 51 [46-61] | 36 [24-49] |
| Eosinophils (%) | 0.25 [0-1.75] | 0 [0-0.3] |
| Macrophages (%) | 42 [30-46] | 55 [32-69] |
| Lymphocytes (%) | 1.5 [0.3-2.0] | 1.1 [0.8-2.0] |
| Bronchial epithelial cells (%) | 2.5 [1.0-8.3] | 2.1 [1.5-3.6] |
| Data are expressed as median [interquartile range] |  |  |

<sup>§</sup>p < 0.05 compared with swimmers; <sup>†</sup>p < 0.05 compared with winter sport athletes.

**Figure 2.**
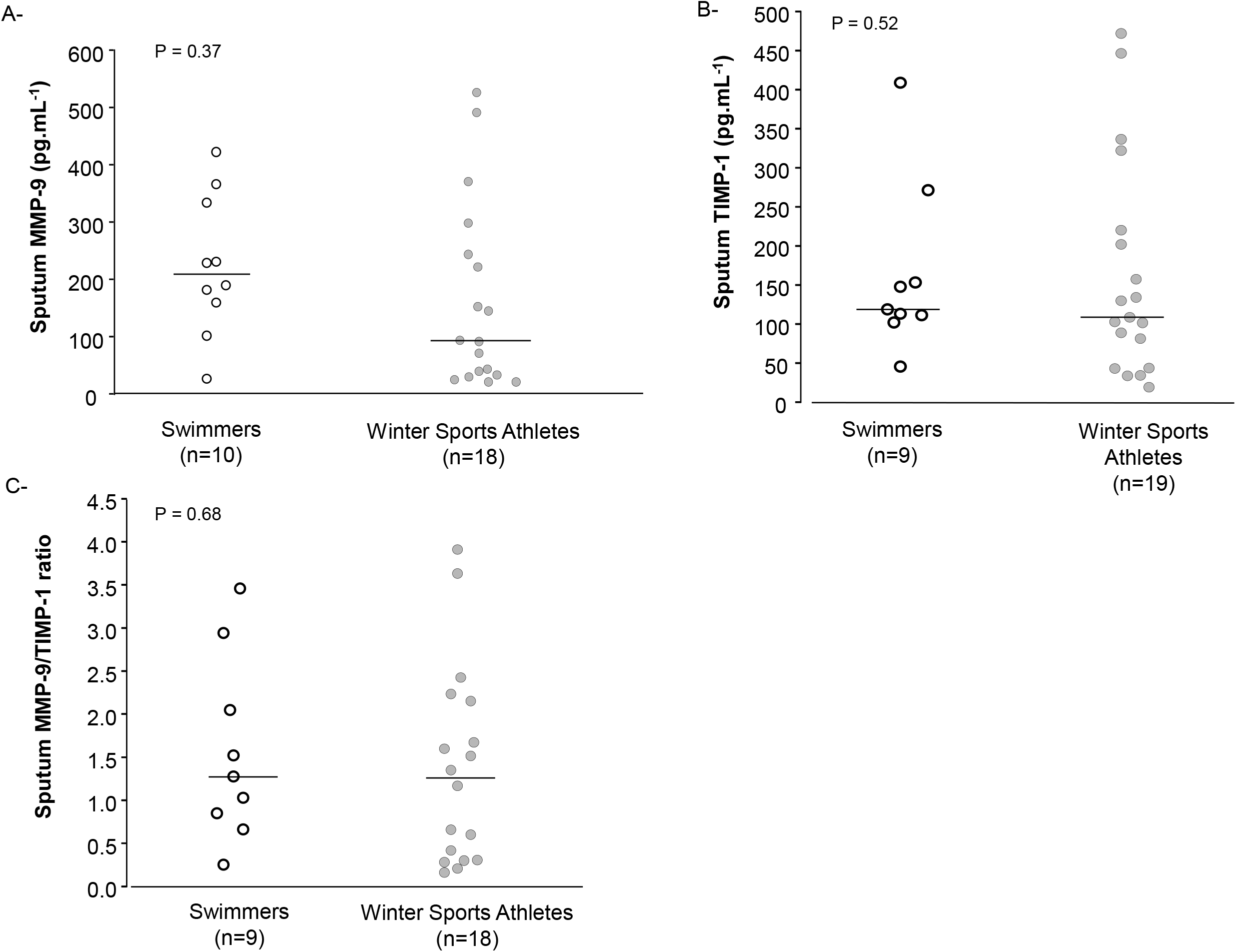
Values of inflammatory cells in induced sputum of athletes. MMP-9: Metalloproteinase-9; TIMP-1: Tissue inhibitor of metalloproteinase-1. The horizontal bar represents the median for each group.

No significant correlation was observed between MMP-9, TIMP-1, MMP-9/TIMP-1 in serum or sputum and training history, training hours /week, FEV_1_, FVC, MIT PC_20_, the maximum fall in FEV_1_ post-EVH, AHR status to MIT and EVH, serum CRP or fibrinogen, serum cc16/SP-D and sputum inflammatory cells in the whole athletes group, in swimmers or in winter sport athletes only. Sputum MMP-9 and TIMP-1 correlated positively in the whole group of athletes (r_s_=0.47, p=0.013), but no correlation was observed between serum MMP-9 and TIMP-1 values (p=0.96). Sputum MMP-9 in athletes (n=27) correlated to serum cc16/SP-D ratio (r_s_=0.45, p=0.019), and MIT PC_20_ (r_s_=-0.47, p=0.013) (Figure 3).

**Figure 3.**
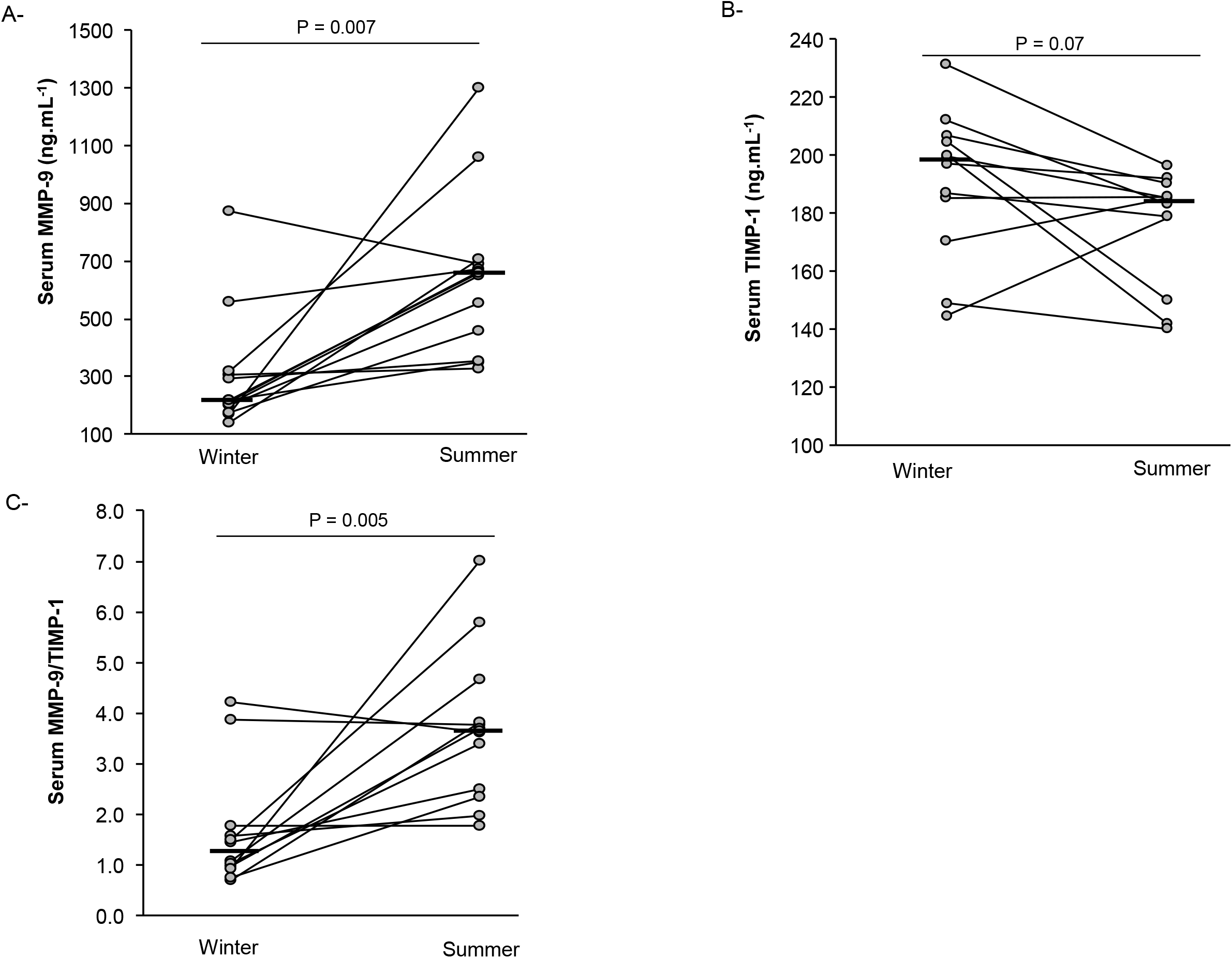
Correlation between sputum MMP-9 and (A) Methacholine airway responsiveness, and (B) serum cc16/SP-D ratio. rs: Spearman correlation

### Comparison of MMP-9 and TIMP-1 during winter and summer seasons in winter sports athletes

Among the 12 winter sports athletes, 11 had atopy, including 9 seasonal atopic. In these athletes, there was a significant increase from the winter to the summer in FEV_1_ (4.67 ± 0.96 L *vs* 4.88 ± 0.95 L, respectively; p = 0.008) and in FVC (5.63 ± 1.13 L *vs* 5.92 ± 1.20 L, respectively; p = 0.003). The difference in FEV_1_ between both seasons was of 207 ± 222 mL (4.29 ± 4.32 %) and of 285 ± 255 mL (4.60 ± 4.07 %) for FVC. Methacholine PC_20_ was not significantly different from the winter to the summer (p=0.30). The FEV_1_ fall in EVH was significantly higher during the summer compared to winter (10.5% [7.2 -14] *vs* 7.6% [4.8 – 11.3], respectively; p = 0.002). The minimum FEV_1_ post-EVH was not significantly different between both seasons (4.36 ± 0.93 L *vs* 4.30 ± 0.93 L, in winter and in summer, respectively; p = 0.72).

Serum MMP-9 and MMP-9/TIMP-1 ratio were significantly greater, whereas serum TIMP-1 trend to be lower, during the summer (Figure 4). Serum fibrinogen was significantly lower during the summer (2.84 ± 0.35 g.L^-1^ in winter *vs* 2.50 ± 0.35 g.L^-1^ in summer; p = 0.04), but no significant difference was observed in serum CRP (p = 0.87). No significant correlation was observed between changes from summer to the winter of serum MMP-9, TIMP-1, MMP-9/TIMP-1 ratio and the changes of FEV_1_, FVC, MIT PC_20_ or FEV_1_ fall post-EVH.

**Figure 4.**
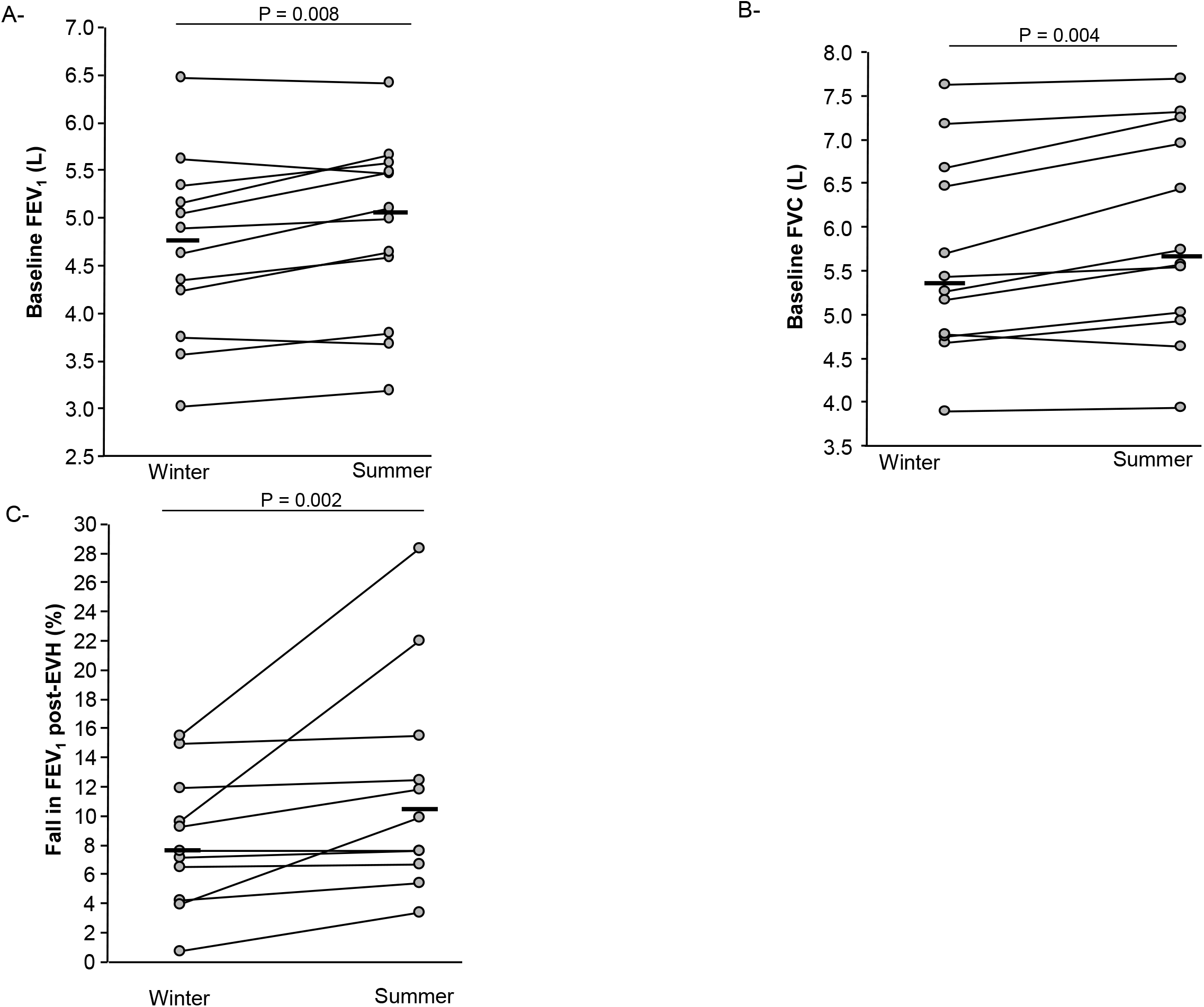
Serum values of inflammation/remodeling biomarkers in winter sport athletes during the winter and the summer. MMP-9: Metalloproteinase-9; TIMP-1: Tissue inhibitor of metalloproteinase-1. The horizontal bar represents the median for each season. CC16: Club cell proteins; MMP-9: Metalloproteinase-9; Methacholine PC_20_: concentration of methacholine inducing a 20% fall in FEV_1_; SP-D: surfactant protein-D; rs: Spearman correlation coefficient.

**Figure 5.**
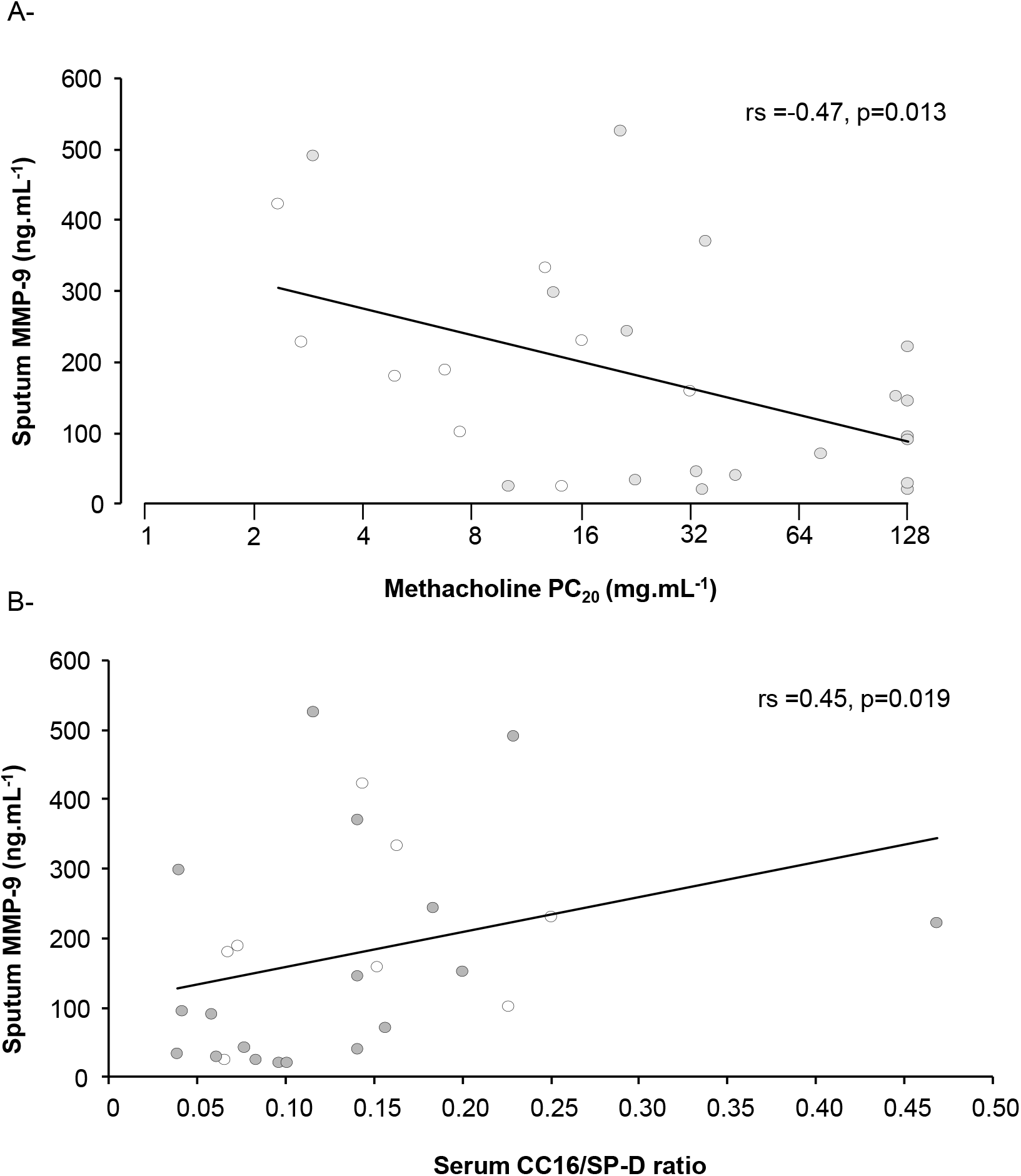

## Discussion

The aims of the study were to compare serum and sputum concentrations of MMP-9 and TIMP-in winter sport athletes and swimmers in relation with airway function at a similar moment of training periodization, and to observe the variations from winter to summer in winter sport athletes. No significant difference in serum or sputum concentrations of MMP-9, MMP-9/TIMP-1 or sputum TIMP-1 was observed between swimmers, winter sport athletes and controls. Sputum MMP-9 correlated with Methacholine PC_20_ and serum cc16/SP-D ratio, suggesting a link between the degree of responsiveness, epithelial damage and MMP-9 in the airways, despite no correlation between cc16/SP-D and MIT PC_20_. Finally, in winter sport athletes, the season of measurement had an impact on serum MMP-9 and TIMP-1, with a higher MMP-9/TIMP-1 ratio observed during the summer, not related to airway function or responsiveness.

The lack of difference in serum MMP-9 and TIMP-1 levels, and MMP-9/TIMP-1 ratio in both athletes’ category and controls, indicates probably that these biomarkers, when measured in blood, are not sensitive nor specific enough in this population to reflect local changes in the airways, that occur after years of training. In that sense, there was no correlation between airway biomarkers of inflammation, responsiveness or function, and serum level of biomarkers. Serum TIMP-1 was moderately increased in swimmers compared with winter sport athletes, probably resulting from a slightly different training recovery state, as TIMP-1 in induced sputum was similar in both groups of athletes. Some studies reported an acute increase in blood MMP-9 after intense exercise, but their origin remains to be determined as studies bring contradictory results (Carmeli *et al*. 2005, Saenz *et al*. 2006, Rullman *et al*. 2007, 2013, Reihmane *et al*. 2012). MMP-9 may originate from several tissues after exercising and participates to the cleaving of muscle-specific proteins and extracellular matrix formation, remodeling and regeneration in skeletal and cardiac muscle, as well as the surrounding vasculature (Koskinen *et al*. 2001, Carmeli *et al*. 2005, Saenz *et al*. 2006, Rullman *et al*. 2013). Blood MMP-9 level may modulate the activation of growth factor cytokines and angiogenesis, thus facilitating physiological adaptations to exercise training. Certain stimuli, especially those producing an important mechanical stress, activate the local production of MMP-9 in the skeletal muscle (Carmeli *et al*. 2005, Rullman *et al*. 2007), and probably also in the airways. The slight increase in serum TIMP-1 unlikely originates from the airways but may reflect an incomplete tissue repair process in the swimmers, due to an insufficient time to recover from the last training and/or chlorine exposure. We chose the same season to compare serum from swimmers and winter sport athletes as it is their annual period of training resumption. However, the new swimming training year, that was also an Olympic year for summer sports, began with a high volume of training, compared with the beginning of the winter sport athletes’ season. In the summer, in winter sport athletes, the serum MMP-9/TIMP-1 ratio was increased whereas the serum TIMP-1 and fibrinogen were decreased compared to the winter season. This is in favour of an increase inflammatory process during the summer, and fibrogenesis in winter. This is an interesting observation meaning that these biomarkers are sensitive to training but also probably to the environment. As all, except 3, had seasonal atopy, we can not exclude a role of allergies in the increase of serum MMP-9.

In acute inflammation due to ozone inhalation, MMP-9 may play a protective role against inflammation in limiting neutrophils’ recruitment and protecting from epithelial injuries and increases in permeability (Yoon *et al*. 2007). MMP-9 have also been shown to promote the migration of repairing bronchial epithelial cells after injury (Legrand *et al*. 1999). As Vermeer *et al*. (2009) observed *in vitro* a loss of bronchial epithelium integrity through the cleavage of one or more components of tight junctions by MMP-9, the relationship between epithelial damage and MMP-9 remains to be clarified. In our study, sputum MMP-9 correlated with serum cc16/SP-D ratio in all the athletes, confirming the link between the loss of epithelial integrity and MMP-9 in the airways, regardless of the sense of the relation between both. Few studies have investigated the relationship between lung function and MMP-9 and results are inconsistent (Vignola *et al*. 1998, Ko *et al*. 2005). In general, in mild stable asthma, there is no link between baseline values of airway MMP-9, that are often similar to the level observed in control subjects, and lung function (Ko *et al*. 2005); but a link is observed between both variables when measured after an allergen challenge (Cataldo *et al*. 2002) or in untreated mild asthmatics having a higher number of neutrophils and macrophages in sputum compared with control subjects (Vignola *et al*. 1998).Todorova **et al**. (2010) suggested that during a stable period of disease, the level of MMP-9 may de determined by their production by structural cells, i.e. bronchial fibroblasts, epithelial cells and smooth muscle cells, whereas during asthma exacerbation, their major production source are inflammatory cells. In their study, the authors reported a positive relationship between the activity of MMP-9 in fibroblasts of stable mild to moderate asthma patients and the dose of methacholine inducing a 20% fall in FEV_1_ (r=0.86, p=0.01). In our study, there is a slight correlation between the level of MMP-9 in sputum and MIT PC_20_ (rs=-0.47, p=0.013), but which is negative. The activity of MMP-9 in the study from Todorova *et al*. (2010) was confined to cultured fibroblasts of asthmatics, whereas in our study they probably originate also from epithelial, and inflammatory cells. Therefore, the level of MMP-9 may reflect an underlying inflammatory cells activity, and a slight damage in epithelial integrity. In that sense, sputum MMP-9 also correlated to serum cc16/SP-D ratio.

A limitation is that we had insufficient induced sputum to analyze in winter sport athletes, and, therefore, we had no measurement of MMP-9 and TIMP-1 in sputum during winter and summer. This may have help us to better understand the origin of MMP-9 during both seasons. However, as induced sputum is sometimes difficult to obtain in athletes (Bougault *et al*. 2009), we thought serum biomarkers as an attractive alternative. However, we found no correlation between serum and sputum values. As all, except 3, had seasonal atopy, we can not exclude a role of allergies in the increase of serum MMP-9.

To conclude, we found no major differences in the level of MMP-9 and TIMP-1 in the serum of winter sport athletes and swimmers, except for a slight increase in TIMP-1 in swimmers. This was not correlated to airway responsiveness, lung function, or the level of airway inflammation, indicating probably a more systemic origin of these markers, related to training and environment. Sputum level of MMP-9 correlated to serum CC16/DP-D ratio and Methacholine PC_20_, suggesting a probably indirect link between the degree of responsiveness, epithelial damage and MMP-9 in the airways. Finally, as serum MMP-9/TIMP-1 ratio was increased in the summer compared with the winter in elite athletes, this may indicate an important role of training and/or environment on these biomarkers, that may also affect the airways. But this remains to be determined. Our study underlines the difficulty to assess inflammation and remodeling mechanisms in athletes’ airways, as serum biomarkers do not reflect the airways’ activity.

## Perspectives

In a near future, priority should be given to search for easy tools and specific protocols to better understand the development of airway structural and functional changes in athletes, according to the training and environmental stimuli. This will make it possible to prevent and better manage respiratory diseases that develop during sports careers, and to identify the phenotypes resulting in identical symptoms.

